# Genomic and phenotypic characterization of *Burkholderia* isolates from the potable water system of the International Space Station

**DOI:** 10.1101/753087

**Authors:** Aubrie O’Rourke, Michael D. Lee, William C. Nierman, Chris L. Dupont

## Abstract

The opportunistic pathogens *Burkholderia cepacia* and *Burkholderia contaminans*, both genomovars of the *Burkholderia cepacia* complex (BCC), are frequently cultured from the potable water system (PWS) of the International Space Station (ISS). Here, we sequenced the genomes and conducted phenotypic assays to characterize these *Burkholderia* isolates. All recovered isolates of the two species fall within monophyletic clades based on phylogenomic trees of conserved single-copy core genes. Within species, the ISS PWS strains all demonstrate greater than 99% average nucleotide identity (ANI), suggesting that they are of a highly similar genomic lineage and both individually may have stemmed from the two founding clonal strains before diverging into two unique sub strain populations. No evidence for horizontal gene transfer between the populations was observed. Differences between the recovered isolates can be observed at the pangenomic level, particularly within putative plasmids identified within the *B. cepacia* group. Phenotypically, the ISS-derived *B. cepacia* isolates generally exhibit a trend of lower rates and shorter duration of macrophage intracellularization compared to the selected terrestrial reference strain (though not significantly), and significantly lower rates of cellular lysis in 7 of the 19 isolates. ISS-derived *B. contaminans* isolates displayed no difference in rates of macrophage intracellularization compared to the selected reference, though generally increased rates lysis, with 2 of the 5 significantly increased at 12-hours post inoculation. We additionally find that ISS-isolated *B. contaminans* display hemolytic activity at 37°C not demonstrated by the terrestrial control, and greater antifungal capacity in the more recently collected isolates. Thankfully, the ISS-derived isolates generally exhibit 1-4 times greater sensitivity to common antibiotics used in their clinical treatments. Thus, despite their infection potential, therapeutic treatment should still have efficacy.

**Author Summary:** The International Space Station (ISS) is a unique built environment due to its isolation and recycling of air and water. Both microbes and astronauts inhabit the ISS, and the potential pathogenicity of the former is of great concern for the safety of the latter. The potable water dispenser (PWD) of the potable water system (PWS) on board the ISS was assembled in a cleanroom facility and then primed on Earth using an extensive process to ensure no gas bubbles existed within the lines that could lock the apparatus upon installation in orbit. The primed system sat dormant for 6 months before installation on the ISS. Microbial surveillance was conducted on the system after installation and the bacterial load was 85 CFU/mL, which exceeded the 50 CFU/mL limits set for ISS potable water. Over a microbial surveillance spanning 4.5 years, numerous strains of the potential pathogen *Burkholderia* have been isolated from the PWD. Here we sequenced and analyzed the genomes of these strains while also characterizing their potential pathogenicity. The genome analysis indicates it is likely that there were only two strains that were introduced on Earth that have subsequently undergone minimal diverging evolution. These strains retain pathogenicity, but remain susceptible to antibiotics, providing a potential therapeutic intervention in the event of infection.

## Introduction

Microbial surveillance of the surfaces, air, and potable water system (PWS) of the International ISS has been implemented by National Aeronautics and Space Administration (NASA) to ensure crew health within this unique closed environment. These efforts, which use standard culturing techniques, have been conducted over twelve years and 22 missions and began shortly after the potable water dispenser (PWD) was launched on STS-126 in November of 2008. On-orbit operations using the PWD began in early 2009 and continues operations to this day. The organisms *Burkholderia cepacia* and *B. contaminans*, both genomovars of the *Burkholderia cepacia* complex (BCC), are frequently cultured from the PWD of the ISS. Our isolates were collected between January 6, 2010 during mission 22 to August 6, 2014 during mission 40 (Fig. 1, S1 Table). The PWS in combination with the PWD is a water recycling system that utilizes physical and chemical techniques to filter, decontaminate, and sterilize water used for drinking and food hydration [1–3]. *Burkholderia* spp. are known to withstand disinfection and sterilization procedures as they display a moderate to high-tolerance to stress such as UV-C radiation, antibiotics, and high heavy-metal concentrations [4].

**Fig 1:**
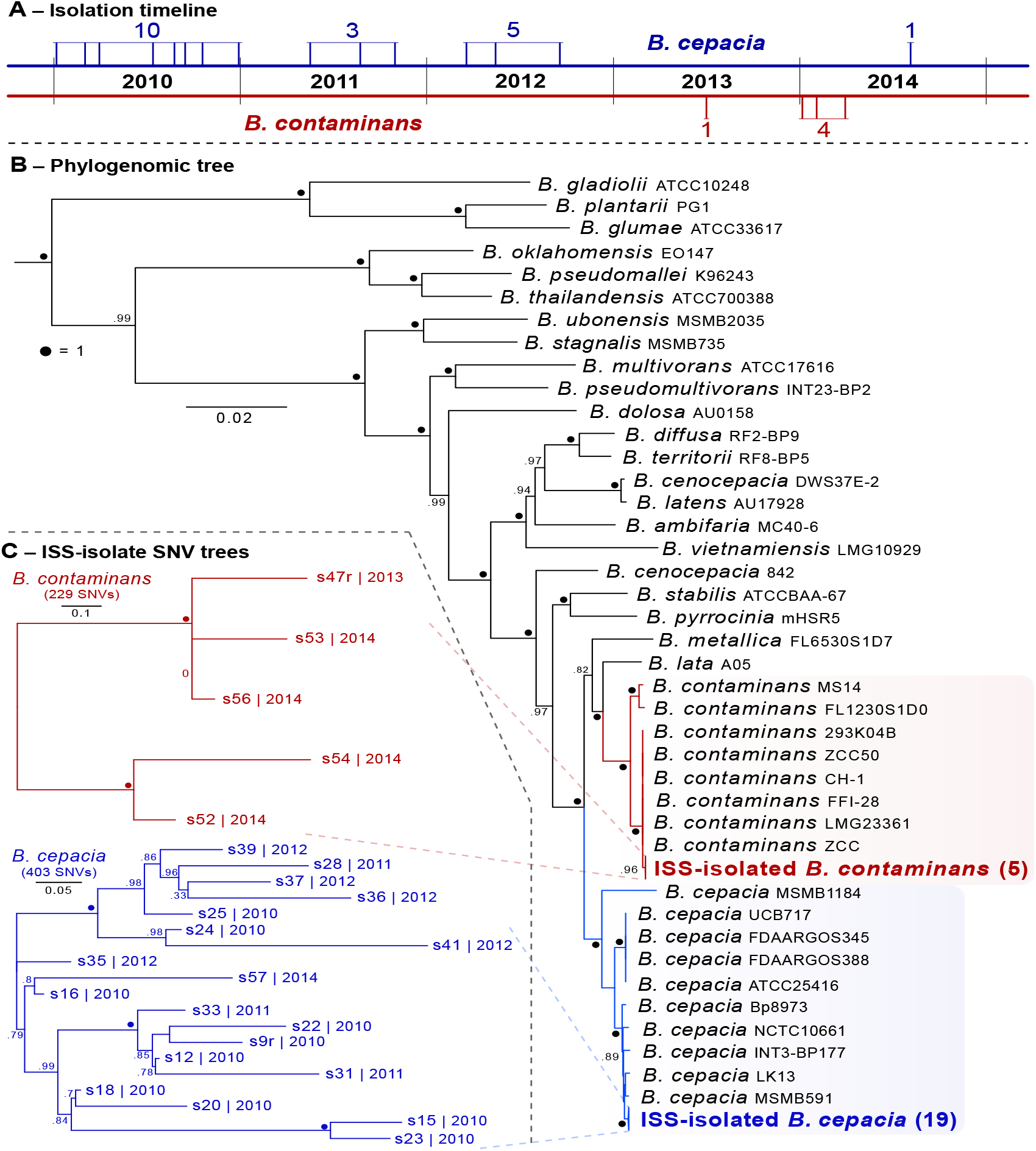
Isolate phylogenetics. **(A)** Isolation timeline (dates with multiple isolates are represented by a single line). **(B)** Estimated maximum-likelihood phylogenomic tree based on aligned and concatenated amino acid sequences of 203 single-copy genes designed for targeting Betaproteobacteria – rooted with *Ralstonia pickettii* 12J (GCF_000020205.1). Numbers in parentheses indicate number of isolates in that clade. **(C)** ISS-derived isolate SNV trees for each *Burkholderia* species. Numbers following the identifiers are the year of isolation for that isolate.

*Burkholderia* spp., among other bacterial contaminants, are likely to have been introduced into the PWD during unit assembly on planet [1–3]. *Burkholderia* spp. probably sustained in the system due to the ability of this genera to survive long periods in distilled water [5,6]. This ability to survive in distilled water with minimal additives has made them problematic for healthcare, as hospital-acquired BCC infections can arise from contaminated disinfectants, anesthetic solutions, distilled water, and aqueous chlorhexidine solutions [4]. The ISS PWD uses flushes of 40ppm elemental iodine for decontamination, followed by iodine removal by a deiodination filter before astronaut use [1–3]. Some *Burkholderia* isolates have been found to be resistant to iodine and can even survive in iodine solutions [7] providing pre-observed phenotypes in this lineage capable of continued presence in the ISS PWD.

*Burkholderia* species are known to propagate in both nutrient-poor and -rich soil environments, in addition to within human host cells [8]. They are copiotrophs commonly isolated from soil, plant rhizospheres, and water [9], as well as from the sputum of cystic fibrosis patients with chronic infection [10]. BCC members pose a significant threat to individuals with cystic fibrosis (CF; accounting for 85% of all infections in CF patients [10]) and otherwise immune-compromised patients due to an exacerbation of pulmonary infections, which can lead to morbidity and mortality [11]. The antibiotic resistance and intracellular survival capabilities of BCC members make them unamenable to may therapeutics in susceptible patients are exposed to them [12]. The genomes of BCC organisms can mutate rapidly during infections or when subjected to high-stress conditions [13], the latter of which would be satisfied by the sequential iodine treatments of the PWD. The genus is known to have several mobile genetic elements (MGEs), which can promote the transfer and acquisition of virulence and antibiotic resistance genes; BCC organisms have genomic islands, such as the *B. cepacia* epidemic strain marker (BCESM), containing genes linked to virulence and metabolism, quorum sensing, transcriptional regulation, fatty acid biosynthesis, and transposition [13]. Here we characterize genomic and phenotypic properties of 24 *Burkholderia* isolates derived from the ISS PWD.

## Results

The PWD unit of the ISS was assembled in a cleanroom facility and then primed on Earth using an extensive process to ensure no gas bubbles existed within the lines that could lock the apparatus upon installation in orbit. The primed system sat dormant for 6 months before installation on the ISS [1,3]. Microbial surveillance was conducted on the system after installation and the bacterial load was 85 CFU/mL, which exceeded the 50 CFU/mL limits set for ISS potable water, leaving the sole source of water on the US module out-of-order [1]. In the meantime, the Russian system was used as a back-up. The US system was flushed with the biocide iodine (I_2_), first at what turned out to be a sub-inhibitory concentration of 4ppm, as subsequent measurements revealed an increase in the microbial load [1]. Further testing revealed that 40ppm was the necessary concentration of iodine flush to achieve the drinkable 50 CFU/mL bacterial load [1]. Iodine flushes are still intermittently administered to the system after durations of PWS stagnation. Our focal *Burkholderia* species are known to survive not only in distilled water, but also in iodine solutions; therefore, these flushes may have reduced the overall microbial load of the PWS while inadvertently selecting for the *Burkholderia* species within the system. Here we characterized the *Burkholderia* species isolates obtained from the ISS PWS both genomically and phenotypically.

Twenty-four isolates collected over 4.5 years were chosen for sequencing. The genomes of 24 ISS-PWD *Burkholderia* isolates (Fig. 1A) were sequenced and *de novo* assembled with SPAdes v3.12.0 ([14]; assembly summary information in S2 Table). The recovered genomes within each species were all found to have an ANI of greater than 99% with 95–99.9% alignment in all cases (fastANI v1.2; [15]). An estimated maximum-likelihood phylogenetic tree based on amino acid sequences of 203 single-copy genes specific to Betaproteobacteria was generated with GToTree v1.4.4 ([16]; Fig 1B). This placed 19 of the isolates within their own monophyletic clade within the *B. cepacia*, and 5 within their own monophyletic clade within the *B. contaminans* (Fig 1B). Single-nucleotide variant (SNV) trees (Parsnp v.1.2; [17]) generated for each group of isolates revealed no clear groupings based on time of isolation (Fig 1C). The *B. cepacia* were isolated over the entire sampling period, which *B. contaminans* were only isolated during 2013 and 2014.

### Pangenomic Analysis

We first performed a pangenomic analysis of each species incorporating all assemblies available from NCBI (as accessed on 20-Aug-2019). Gene clusters (GCs) were identified with MCL clustering [18] as employed within anvi’o v5.5 [19]. Across the incorporated 155 *B. cepacia* genomes (136 reference from NCBI and 19 derived from the ISS PWS), the average gene count was 7,461 ± 479 (mean ± 1SD). The MCL clustering approach yielded the identification of 25,383 total GCs: 2,589 with genes contributed by all (“core”; 10.2%); 8,249 with genes contributed by only single genomes (“singletons”; 32.5%); and 14,545 contributed by some mixture (“accessory”) (Fig 2A). A total of 25 *B. contaminans* genomes were incorporated (20 references from NCBI and 5 ISS-isolates), with an average gene count of 7,580 ± 539. From these, a total of 13,645 GCs were generated: 3,534 core (25.9%); 3,436 singletons (25.2%); and 6,675 accessory GCs (49.0%; Fig 2B). It should be noted the disparity between the number of identified core, singleton, and accessory genes in *B. cepacia* vs those identified in *B. contaminans* at this point is not meant to convey anything about potential differences in gene-content diversity between the two species. The number of genomes available/incorporated for each and the breadth of phylogenetic diversity spanned by the incorporated genomes of the two groups both vary greatly.

**Fig 2:**
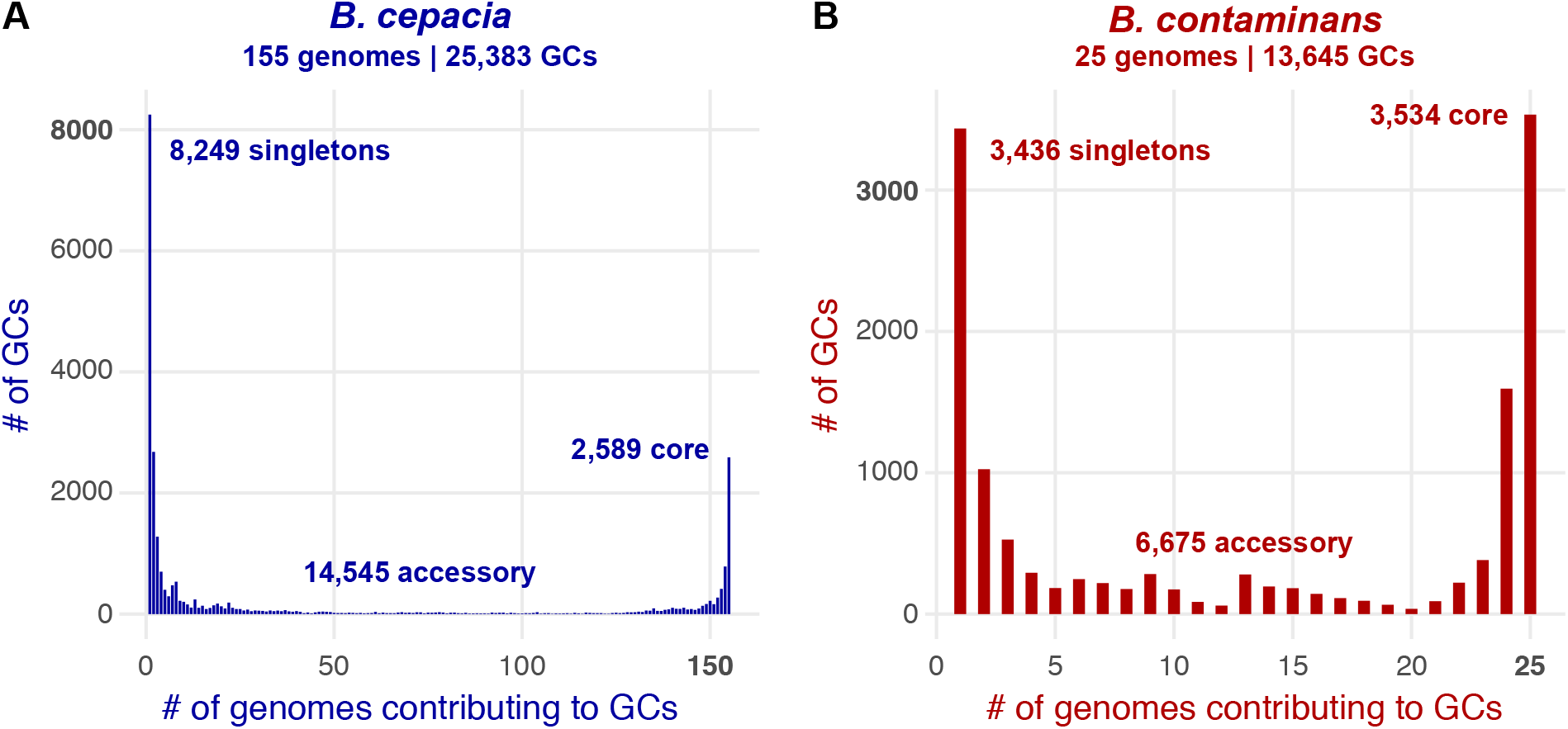
**Gene-cluster distributions** for **(A)** *B*. cepacia and **(B)** *B. contaminans*.

We scanned for functional enrichment or depletion in the ISS-derived isolates as compared to the reference genomes based on normalized frequencies of presence/absence across the two groups (see Methods). It should be kept in mind that given the phylogenetic landscape of both ISS-derived groups – with each forming monophyletic clades within their respective species (Fig 1B) – the functional differences observed may be due to evolutionary divergence as a whole, rather than being due to their source of isolation (the ISS). Of the *B. contaminans* pangenome, this revealed no depleted functions in the ISS-derived isolates, and 8 functions enriched including those commonly associated with viruses and conjugative plasmids (Table 1; full results in S3 Table).

**Table 1:** ISS-derived *B. contaminans* enriched functions as compared to references.

| Annotation (NCBI PGAP) | Occurrence in ISS-isolates | Occurrence in Refs | Corrected p-value |
| --- | --- | --- | --- |
| Invasion protein lagB | 5/5 | 0/20 | 0.09 |
| Nucleoside 2-deoxyribosyltransferase | 5/5 | 0/20 | 0.09 |
| PH domain-containing protein | 5/5 | 1/20 | 0.09 |
| RepA replicase | 5/5 | 1/20 | 0.09 |
| Metallophosphatase family protein | 5/5 | 1/20 | 0.09 |
| Pilus assembly protein PilN | 5/5 | 1/20 | 0.09 |
| Prepilin type IV pili | 5/5 | 1/20 | 0.09 |
| Type IV secretion system protein VirB3 | 5/5 | 1/20 | 0.09 |

In contrasting the functional annotations of the 19 ISS-derived *B. cepacia* with the 136 references, 265 functions were found to be enriched, with 98 found to be depleted (5 of each presented in Table 2; full results in S4 Table).

**Table 2:** ISS-derived *B. cepacia* enriched/depleted functions as compared to references.

| Annotation (NCBI PGAP) | Occurrence in ISS-isolates | Occurrence in Refs | Corrected p-value |
| --- | --- | --- | --- |
| Toprim domain-containing protein | 19/19 | 0/136 | 0.00 |
| 3-carboxyethylcatechol 2,3-dioxygenase | 19/19 | 0/136 | 0.00 |
| Metal transporter | 19/19 | 1/136 | 0.00 |
| Plasmid mobilization relaxosome MobC | 19/19 | 1/136 | 0.00 |
| Peptidase M4 | 19/19 | 3/136 | 0.00 |
| Sugar transporter | 0/19 | 133/136 | 0.00 |
| LTA synthase family protein | 0/19 | 132/136 | 0.00 |
| Class II aldolase family protein | 0/19 | 123/136 | 0.00 |
| Nit6803 family nitriliase | 0/19 | 111/136 | 0.00 |
| MSMEG_0572 N starvation response | 0/19 | 111/136 | 0.00 |

As noted above, the recovered genomes within each species were all found to have an ANI of greater than 99% with 95–99.9% alignment in all cases, and each form monophyletic clades in relation to other members of their respective species (Fig 1B). Pangenomic analyses of solely the ISS-derived isolates revealed highly conserved cores of GCs for both species, with greater than 90% of the GCs having genes contributed from all genomes. For the 19 *B. cepacia* isolates, 7,688 total GCs were generated: 7,065 core (91.9%); only 20 singletons (0.26%); and 603 accessory GCs (7.8%; Fig 3, left side). Of the singleton GCs with functional annotations (5/20), most annotations were present in other GCs – meaning the sequences diverged enough to not cluster together, but were similar enough to be annotated the same way. The one exception was annotated by NCBI as an autotransporter domain-containing protein (from isolate s20, gene ID D7204_40525). For the 5 *B. contaminans* isolates, 7,778 GCs were generated: 7,509 core (96.5%); 163 singletons (2.1%); and 106 accessory GCs (1.4%). Of the 163 singleton GCs from *B. contaminans*, 46 had annotations associated with them. Twenty-eight of these annotations were found in other GCs, while 18 were only annotated in one genome. Interestingly, 17 of these 18 all came from the same isolate s47, which was the first *B. contaminans* isolate recovered from the ISS (Fig 1C); these functional annotations are presented in Table 3. All gene calls, sequences, and annotations are available in S5 Table.

**Fig 3:**
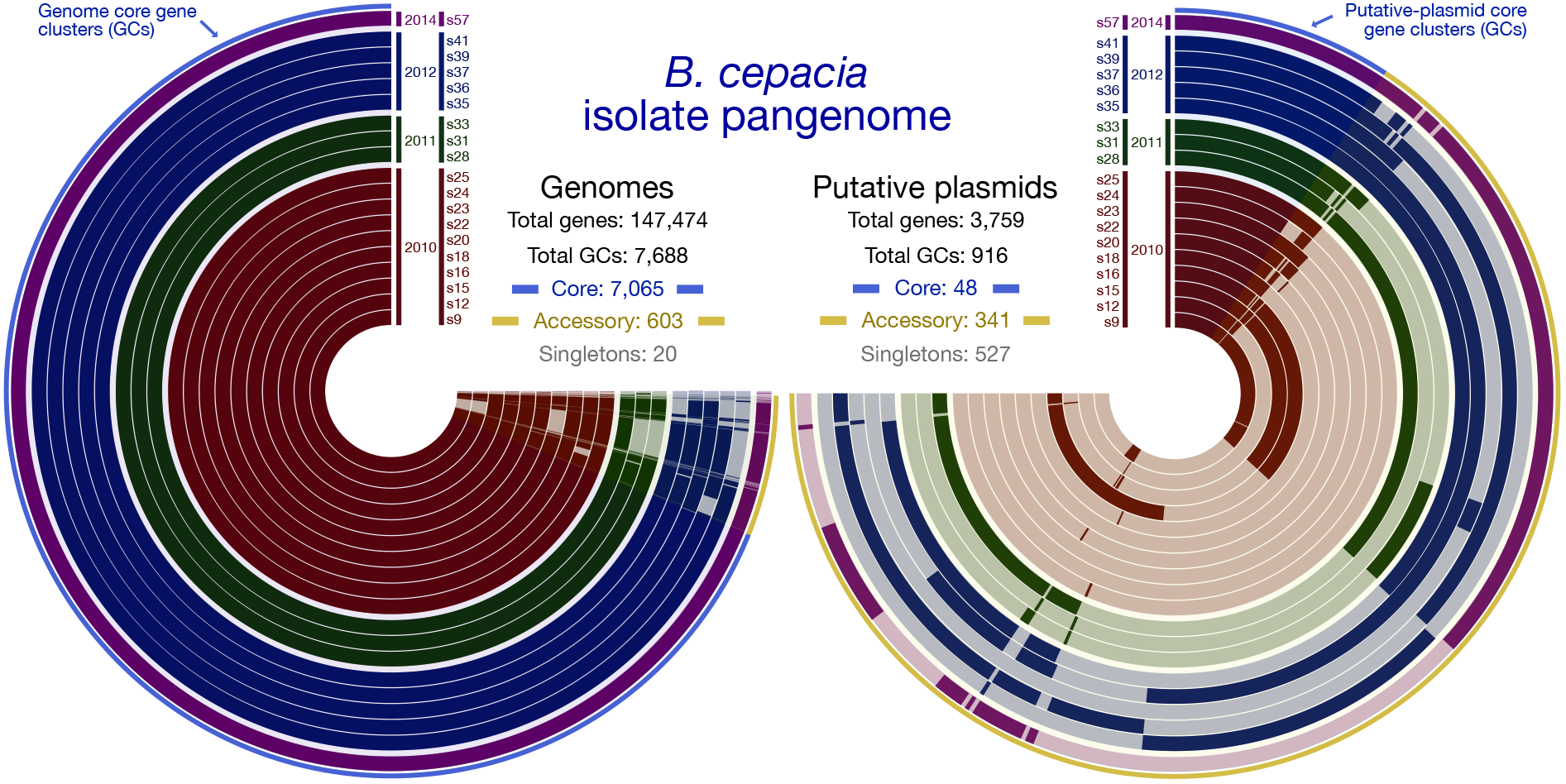
Pangenomics visualizations of the 19 ISS B. cepacia isolate genome assemblies (left) and identified putative plasmids (right). Each concentric circle radiating out from the center represents an isolate, identified at the top of each next to the year they were isolated. Wrapping around the circles are the generated gene clusters (GCs), where a solid mark for an isolate at a given GC indicates that particular isolate contributed a gene to that gene cluster, and the absence of a solid color indicates that isolate did not contribute a gene to that gene cluster. The very outer layer of each is a blue and yellow line. The blue line highlights core gene clusters. The yellow line highlights accessory GCs. The putative-plasmid pangenome on the right does not include singletons in the visualization.

**Table 3:** *B. contaminans* singleton-GC functional annotations not identified in any other GCs.

| Annotation (NCBI PGAP) | Isolate | Gene ID |
| --- | --- | --- |
| Nickel resistance protein | s47 | D7S92_31485 |
| Conjugal transfer protein TrbE | s47 | D7S92_31155 |
| Conjugal transfer protein TrbI | s47 | D7S92_31155 |
| DUF1259 domain-containing protein | s47 | D7S92_31415 |
| C-type cytochrome biogenesis protein CcsB (ccsB) | s47 | D7S92_31465 |
| Type IV secretion system family protein | s47 | D7S92_31160 |
| Toprim domain-containing protein | s47 | D7S92_35590 |
| P-type DNA transfer ATPase VirB11 (virB11) | s47 | D7S92_31190 |
| Replication initiator protein A | s47 | D7S92_31330 |
| Type II secretory pathway component PulF-like protein | s47 | D7S92_31550 |
| Chromate resistance protein | s47 | D7S92_31420 |
| Type IV secretion system protein VirD4 | s47 | D7S92_31125 |
| Type IV secretion system protein VirB8 | s47 | D7S92_31175 |
| Zinc metalloproteinase Mpr protein | s47 | D7S92_31335 |
| Zeta toxin family protein | s47 | D7S92_31320 |
| Nickel/cobalt efflux protein RcnA | s47 | D7S92_31490 |
| Conjugative relaxase | s47 | D7S92_31130 |
| Type IV secretion protein Rhs | s56 | D7S91_34560 |

### Plasmid Analysis

Putative plasmids were computationally identified in the ISS-derived isolates using plasmidSPAdes [20], which operates largely based on coverage. This process identified putative plasmid contigs from each of the 19 *B. cepacia* isolates and 2 of the 5 *B. contaminans* isolates (s47 and s52; annotations and sequences for all can be found in S6 Table. Running a pangenomic analysis on the 19 *B. cepacia* putative plasmids generated a total of 916 GCs: 48 core (5.2%); 527 singletons (57.5%); and 341 accessory GCs (37.2%; Fig 3, right side). Despite the high similarity between the ISS genomes as a whole, the majority of variability that does exist within their genetic complement (Fig 3, left side ≈3-3:30 “o’clock”) appears to be sustained among the putative plasmids that were identified (Fig 4, right side). The 2 recovered putative plasmids from *B. contaminans* had little overlap based on GCs. With a total of 475 GCs, only 3 had genes contributed from both genomes (0.6%), leaving the remaining 472 as singletons. Annotations from NCBI’s Clusters of Orthologous Genes (GOGs; [21])of coding sequences from the putative plasmids reveal elements typical of conjugative plasmids, such as DNA replication proteins (DnaC), plasmid stabilization proteins (ParE), and Type IV secretion system components (T4SS; Table 4). Additionally, in *B. cepacia* putative plasmids were identified a catechol 2,3-dioxygenase, a contamination indicator as it is able to breakdown polycyclic aromatic hydrocarbons [22,23], and lysophospholipase, an enzyme known to be used by ingested bacteria to avoid phagocytosis by macrophage [24]. Despite containing this small conserved core, the larger plasmids in particular contain multiple copies of genes with annotations such as bacteriophage DNA transposition protein, AAA+ family ATPase (s9, s36, s39, s57), Virulence-associated protein-VagC (virulence associated gene C; s16, s28, s35, s36, s39), and additional elements of the T4SS, VirB8 (S6 Table). We additionally see transposases-related genes in the *B. cepacia* isolates with a mean copy number per putative plasmid of 17.57 ± 5.7 (mean ± 1 SD), and an integrase element with a mean copy number of 2.47 ± 0.51, suggesting DNA rearrangements and duplications among the conjugative plasmids of ISS *B. cepacia* may be a result of these mobile genetic elements.

**Fig 4:**
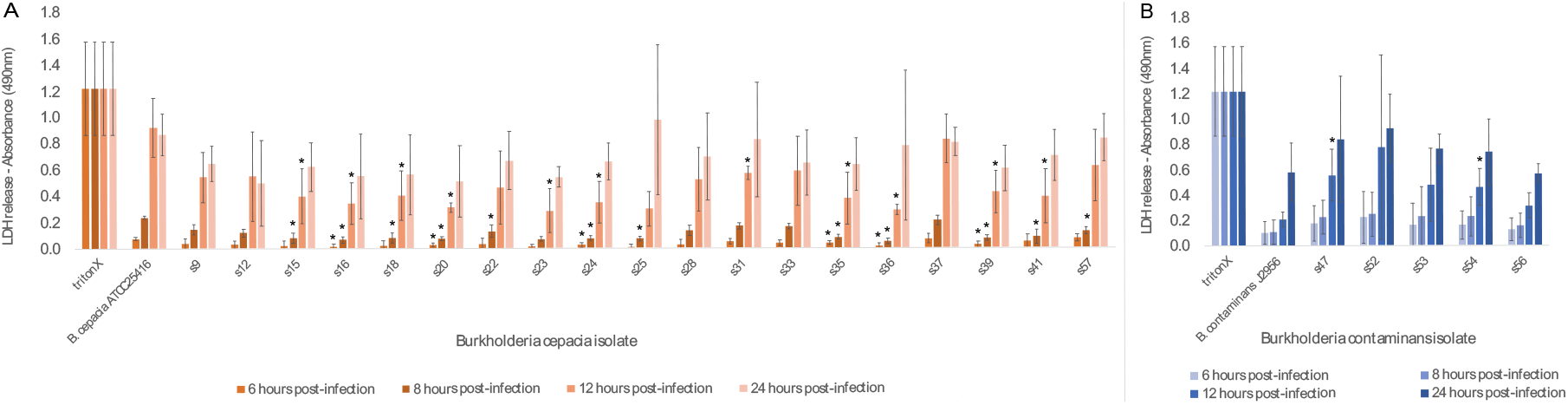
**Lactase dehydrogenase (LDH) release from macrophage** 8 hours, 12 hours and 24 hours post exposure to Burkholderia isolates **(A)** *B.cepacia* and **(B)** *B. contaminans* along with terrestrial reference strains. Increase absorbance correlates to increase LDH release from macrophage. Symbols indidcate significant difference from corresponding terrestrial reference at 0.05 threshold. “TritonX” exemplifies complete lysis.

**Table 4:** COG functional annotations and copy numbers per putative plasmid for *B. cepacia* and *B. contaminans*.

| *B. cepacia* putative-plasmid annotations (19) |  |
| --- | --- |
| Annotation (COG) | Mean copy # ± 1 SD |
| Plasmid stabilization system protein ParE | 1 ± 0 |
| Site-specific DNA-cytosine methylase | 1 ± 0 |
| Type IV secretory pathway, component VirB8 | 1 ± 0 |
| Lysophospholipase | 1.11 ± 0.5 |
| Catechol 2,3-dioxygenase | 1.84 ± 1.1 |
| Arsenite efflux pump ArsB, ACR3 family | 1.89 ± 1.1 |
| Integrase | 2.47 ± 0.5 |
| DNA replication protein DnaC | 3.05 ± 0.2 |
| Cu/Ag efflux pump CusA | 3.05 ± 3.4 |
| DNA-binding transcriptional regulator, LysR family | 3.11 ± 5.9 |
| Transposase-related | 17.11 ± 5.7 |
| *B. contaminans* putative-plasmid annotations (2) |  |
| Annotation (COG) | Mean copy # ± 1 SD |
| Cellulose biosynthesis protein BcsQ | 1 ± 0 |
| mRNA-degrading endonuclease, toxin component of the MazEF toxin-antitoxin module | 1 ± 0 |
| Tfp pilus assembly protein PilT, pilus retraction ATPase | 1 ± 0 |
| Type II secretory pathway component GspD/PulD (secretin) | 1.5 ± 2.1 |
| Type IV secretory pathway, component VirB8 | 1.5 ± 2.1 |
| Type IV secretory pathway, VirB9 components | $2 \pm 1.4$ |
| Chromosome segregation protein ParB | $2 \pm 0$ |
| Chromate transport protein ChrA | $2 \pm 0$ |
| DNA-binding protein H-NS | $2.5 \pm 0.7$ |
| Soluble lytic murein transglycosylase | $3 \pm 2.8$ |
| Transposase-related | $5 \pm 5.7$ |

As for *B. contaminans*, in addition to harboring elements of both the T4SS and Type II secretion system (T2SS), the plasmids have a soluble lytic murein transglycosylase harboring a putative invasion domain LysM at a mean copy number of 3 ± 2.8. The lytic transglycosylase, LtgG, has recently reported for its role to control cell morphology and virulence in *Burkholderia pseudomallei* [25]. Furthermore, we see each harboring a copy of the toxin component of the MazEF toxin-antitoxin system suggesting these ISS *B. contaminans* isolates may be able to control their transition to the dormant persister state [26]. Again, we see transposase-related genes with a mean copy number of 5 ± 5.7.

### Macrophage Infection Assay

The identification of putative virulence mechanisms within the conjugative plasmids of the ISS *Burkholderia* genomes led us to explore the ability for the ISS *B. cepacia* and *B. contaminans* isolates to invade and persist within macrophage in comparison to the terrestrial reference strains of *B. cepacia* ATCC25416 and *B. contaminans* J2956 in a cell culture. For reference, we have included two terrestrial strains from the same lineages (*B. cepacia* and *B. contaminans*), however must stress that these are not controls as they are different strains. Ideal controls would be terrestrial strains from within the monophyletic clades of ISS isolates. We used two metrics for this assessment: 1) the ability of the bacteria to invade macrophage cells and be retained intracellularly, which allows for the bacteria to replicate without lysosomal degradation in turn lending to chronic infections; and 2) the ability for the bacteria to cause macrophage cell lysis as assessed by the amount of lactate dehydrogenase (LDH) released by the cell. Such cell lysis in turn can trigger cytokine release and an inflammatory response. We present the cell counts for the intracellularized bacteria and LDH data for all ISS-derived isolates and *B. contaminans* J2956 and *B. cepacia* ATCC25416 reference strains. Macrophage cell lysis was quantified by the amount of LDH released in the cell culture media at six, eight, twelve and 24 hours post-inoculation (Fig 4). Bacteria which were internalized by macrophage were quantified and reported in colony forming units per milliliter (CFU/mL) measured at six and twelve hours post-inoculation (Fig 5).

**Fig 5:**
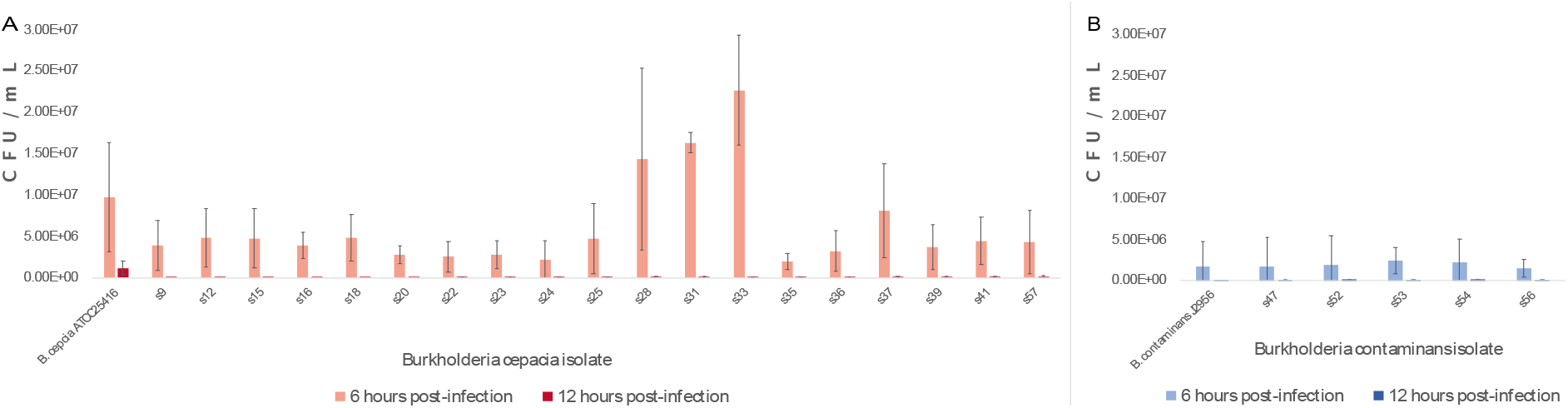
**Colony forming unit per milliliter (CFU/ml) counts** of Burkholderia isolates **(A)** *B.cepacia* and **(B)** *B. contaminans* and terrestrial reference strain intracellularized by J774A.1 mouse macrophage.

At six and eight hours post infection, little LDH-release was observed for both the ISS-derived *B.cepacia* isolates and terrestrial control strain (Fig 4A). The *B. contaminans* ISS-derived isolates appear to be triggering cell lysis at a slightly greater rate than the terrestrial control, though with high variability across the experimental triplicates (Fig 4B) – likely due to higher rates of macrophage lysis observed in experiment two (S2 Fig). At six and eight hours, ISS isolates of *B. contaminans* in general display greater cellular lysis rates than those of the either the *B. cepacia* ATCC25416 or *B. contaminans* J2956 terrestrial control strains, though not significantly. At twelve hours post-infection, the lysis trend exhibited for each isolate exists but at twice the magnitude of LDH released at eight hours post infection (with s47 and s54 significantly greater than the terrestrial control J2956). Finally, at 24 hours post-infection, LDH release approaches that of triton-X, an example of full lysis (Fig 4).

At twelve hours post-infection it becomes apparent that the terrestrial strain of *B. cepacia* ATCC25416 is able to survive intracellularly within macrophage to a greater degree than any of the ISS isolates (Fig 5). It is this ability to persist within macrophage that tends to lead to chronic infection.

### Antifungal and Hemolysis Assay

Non-ribosomal peptide synthetase (NRPS)-derived occidiofungin/burkholdine-like compounds are produced by *B. contaminans* via a gene cluster thought to have evolved to protect BCC bacteria from ecological niche predators such as amoeba and fungi [27]. Using antiSMASH [28] we verified that each of the ISS isolates and the reference strain contain the occidiofungin and pyrrolnitrin gene cluster. The antifungal metabolite, occidiofungin, as well as pyrrolnitrin are known to also display hemolytic properties [29], which can break down heme in the hemoglobin of host bodies causing complications for the host.

Due to the identification of *B. contaminans* isolates harboring these two biosynthetic gene clusters within our collection, we assayed for the ability to inhibit growth of the *Aspergillus fumigatus* AF293 strain (Fig 6A) and cause hemolysis (Fig 6B). None of the *B. cepacia* isolates exhibited fungal inhibition or hemolysis, but each of the *B. contaminans* did to varying degrees. The terrestrial control reference strain (*B. contaminans* J2956) displayed the least amount of fungal inhibition (Fig 6A) and little to no hemolysis (Fig 6B), while isolate s47 displayed a similar ability to inhibit fungal growth but an added ability to lyse blood cells after 48 hours of growth. Though time of isolate-recovery does not necessarily indicate total time that isolate spent aboard the ISS, the isolates that display greater antifungal and hemolytic properties than s47 and the terrestrial control were collected at a later date (Fig 1A and 1C).

**Fig 6:**
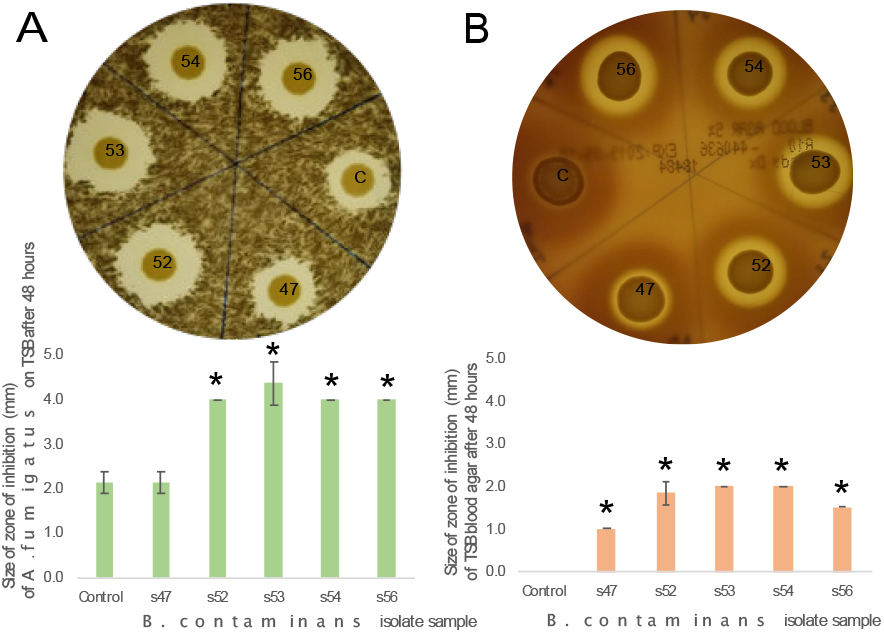
ISS *B. contaminans* – known to produce the antifungal occidiofungin –. **(A)** zones of inhibition formed on Aspergillus fumigatus. **(B)** Zones of inhibition presented by ISS *B. contaminans* on TSB plates with 5% sheep’s blood.

### Minimum Inhibition Concentration Assay

ISS isolates and terrestrial reference control isolates were assayed for antibiotics commonly used in the clinical treatment of infections they are responsible for. These antibiotics included: cefotaxime, meropenem, and ceftazidime (cell wall synthesis inhibitors); ciprofloxacin (a topoisomerase inhibitor); cotrimethoprim (trimethoprim/sulfamethoxazole- a thymidine synthesis inhibitor); chloramphenicol (a 50S ribosomal subunit inhibitor); levofloxacin (a DNA synthesis inhibitor); and minocycline (a 30S ribosomal subunit inhibitor). Most isolates presented a minimum inhibitory concentration (MIC) within 1- to 4-fold that of the terrestrial control strains (*B. contaminans* J2956 and *B. cepacia* ATCC25416) for each antibiotic tested (Table 5). S33 was an exception as it appears to be more sensitive to co-trimethoprim. All isolates appeared to be moderately resistant to cefotaxime, yet susceptible to ceftazidime, also a 3^rd^ generation β-lactam cephalosporin.

**Table 5:** Minimum inhibitory concentration (MIC) values for all ISS-derived *Burkholderia* isolates and incorporated refence control strains.

| ug/mL | ceftazidime | cefotaxime | chloramphenicol | ciprofloxacin | cotrimethoprim | levofloxacin | minocycline | meropenem |
| --- | --- | --- | --- | --- | --- | --- | --- | --- |
| B. cepacia<br>ATCC25416 | 12 | 64 | 6 | 1 | 24 | 1 | 3 | 20 |
| s9 | 8 | 64 | 7 | 2 | 17 | 6 | 10 | 32 |
| s12 | 4 | 32 | 6 | 1 | 12 | 4 | 3 | 24 |
| s16 | 4 | 32 | 8 | 1 | 20 | 4 | 5 | 10 |
| s18 | 4 | 32 | 8 | 1 | 20 | 4 | 9 | 18 |
| s20 | 8 | 32 | 8 | 1 | 24 | 4 | 9 | 18 |
| s22 | 16 | 64 | 4 | 1 | 24 | 6 | 6 | 32 |
| s23 | 4 | 64 | 4 | 2 | 24 | 6 | 6 | 16 |
| s24 | 4 | 64 | 6 | 2 | 17 | 6 | 10 | 20 |
| s25 | 8 | 64 | 6 | 1 | 24 | 6 | 6 | 24 |
| s28 | 4 | 64 | 4 | 2 | 24 | 6 | 6 | 20 |
| s31 | 8 | 64 | 4 | 1 | 24 | 4 | 4 | 24 |
| s33 | 4 | 64 | 4 | 1 | 2 | 1 | 2 | 10 |
| s35 | 8 | 64 | 6 | 1 | 48 | 3 | 8 | 32 |
| s36 | 4 | 32 | 18 | 2 | 24 | 1 | 10 | 48 |
| s37 | 8 | 64 | 5 | 2 | 32 | 2 | 6 | 32 |
| s39 | 8 | 64 | 5 | 2 | 24 | 2 | 4 | 48 |
| s40 | 4 | 64 | 24 | 1 | 4 | 2 | 10 | 64 |
| s41 | 8 | 64 | 5 | 1 | 24 | 1 | 10 | 48 |
| s57 | 8 | 64 | 8 | 1 | 16 | 1 | 2 | 6 |
| B. contaminans<br>J2956 | 8 | 64 | 8 | 1 | 8 | 1 | 2 | 4 |
| s47 | 8 | 64 | 6 | 1 | 8 | 1 | 2 | 4 |
| s52 | 8 | 64 | 8 | 1 | 6 | 1 | 2 | 4 |
| s53 | 12 | 64 | 8 | 1 | 8 | 1 | 2 | 4 |
| s54 | 12 | 64 | 8 | 1 | 6 | 1 | 2 | 4 |
| s56 | 8 | 64 | 8 | 1 | 12 | 1 | 3 | 6 |

### Biofilm Assay

The time of collection records show that the *Burkholderia* species are intermittently cultured from the ISS PWD; there are numerous consecutive flights where *Burkholderia* could not be cultivated from the PWD. One reason for this could be due to the proclivity for *Burkholderia* species to form biofilms. This biofilm would be adapted over time to the pressure and flow experienced during water removal from the system as needed for drinking or food hydration. We observed the ability of the *B. cepacia* isolates to form biofilms to a slightly greater degree than the *B. cepacia* terrestrial strains used for comparison (Fig 7A). We find that on a whole, *B. contaminans* form biofilms more readily than *B. cepacia*, yet the ISS isolates of *B. contaminans* have a diminishing ability to form biofilm in relation to the *B. contaminans* reference control strain (Fig 7B).

**Fig 7:**
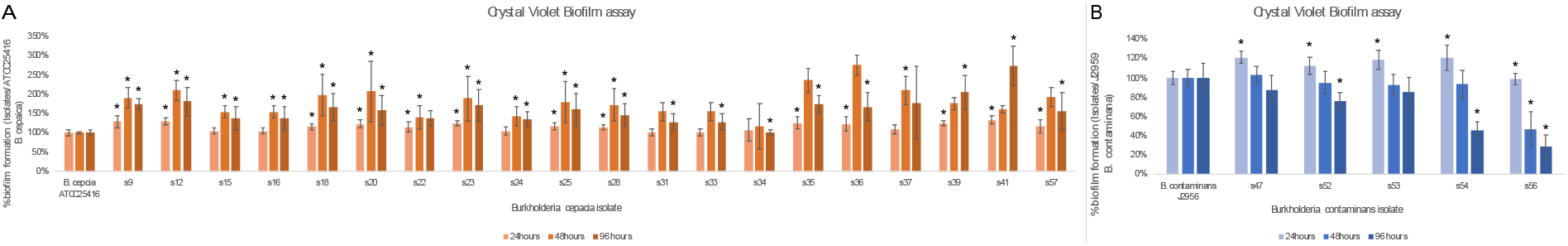
**% biofilm formation** for all ISS *Burkholderia* isolates **(A)** *B.cepacia* and **(B)** *B. contaminans* in relation to the respective terrestrial control strain.

## Discussion

### *Burkholderia* genomic analyses

The ISS-isolated *B. cepacia* and *B. contaminans* both formed monophyletic clades when placed in a phylogenomic tree with currently available references (Fig 1B). When comparing the ISS isolates among one another, they display very few single nucleotide polymorphisms (SNPs) suggesting low genetic diversity among the respective ISS PWD strains. In a pangenomics approach we identified core and accessory GCs for *B. cepacia* and *B. contaminans* and identified the functional annotations enriched and depleted in the ISS-derived isolates as compared to the incorporated references (Tables 1 and 2; S3 Table). There were a small number of functions strictly unique to the ISS isolates. *B. contaminans* isolates each contained two enriched functions annotated as an Invasion protein *iagB* and Nucleoside 2-deoxyribosyltransferase which were not identified in the 20 available terrestrial reference genomes queried. IagB is associated with *Salmonella enterica* subsp. *enterica* ser. Typhi invasion of HeLa cells [30]. *B. cepacia* isolates each contained two enriched functionas annotated as a Toprim domain-containing protein and a 3-carboxyethylcatechol 2,3-dioxygenase that were not found in the 136 terrestrial reference control strains queried. Here, 3-carboxyethylcatechol 2,3-dioxygenase is involved in the breakdown of polycyclic aromatic hydrocarbons (PAHs) to be used as a carbon and energy source [31]. This pangenomic analysis indicates that the ISS isolates are certainly Earth-derived yet they have likely diverged due to the selective pressures imposed by the PWD.

One particular selective pressure that we know of is the iodine disinfection process used to maintain the PWD’s bacterial load, which may have selected for these isolates and further streamlined their genomes to what they are now. Another explanation for their ability to survive these iodine shocks is their displayed propensity to form biofilms which will insulate members of the population from disinfection. Alternatives to iodine have been considered for future PWD microbial disinfection, such as the use of silver in the form of silver (I) fluoride [3,32]. The effect of silver treatment upon *B. cepacia* has been explored as a clinical alternative to antibiotics [33] and will likely reduce this microbial load. However, we find that the ISS *B. cepacia* isolates analyzed in this study all harbored a Cu/Ag efflux pump (CusA) at a mean copy number of 3.05 ± 3.4 on a conjugative plasmid (Table 4). This suggest that they will able to share this ability to pump silver out from within their membrane with the subpopulations of the *B. cepacia* bacterial community that does not yet confer this resistance.

Similarly, we find a number of the identified enriched functions in our pangenomic analysis are commonly associated with mobile genetic elements that can be found on conjugative elements. A more in-depth plasmid analysis reveals that both species harbor elements of the Type IV secretion system (T4SS) on putative plasmids as well as an enrichment of transposase-related genes (S6 Table). Where, T4SSs are multi-subunit cell-envelope spanning structures consisting of a pilus and a secretion channel whereby DNA or protein is translocated outside of the cell to either a target cell or to the surrounding environment. T4SS mediates horizontal gene transfer, which contributes to genomic plasticity and the evolution of pathogens through the spread of antibiotic resistance or virulence genes [34]. The coverage-based plasmidSPAdes [20] approach identified putative plasmids in all 19 of the ISS *B. cepacia* isolates. A pangenomic view of these putative plasmids for *B. cepacia* revealed a conserved functional core of elements including the T4SS VirD4 gene as well as a catechol 2,3-dioxygenase and lysophospholipase (Table 4). Putative plasmids were also identified in 2 of the 5 ISS *B. contaminans* isolates. These held similar functional annotations including lytic transglycosylase and the toxin component of the MazEF toxin-antitoxin system (Table 4).

### *Burkholderia cepacia* plasmid encoded features

The ISS *B. cepacia* isolates all harbor at least one copy of lysophospholipase, this is an enzyme that frees fatty acids from lysophospholipids (LPLs) and in turn generates cytotoxic LPLs. As a result, LPLs are considered to be virulence factors of bacteria as they are found to help bacteria escape phagosomes in host cells after a few rounds of intracellular multiplication [24]. The cytotoxicity they generate allows the bacteria to rupture out of a macrophage or epithelial cell, and in addition, destroy lung surfactant and generate signal transducers such as lysophosphatidylcholine, which in turn can induce inflammation [24].

Catechol 2,3-dioxygenases (C23Os) found on the isolate putative plasmids are commonly used as water quality indicators [35]. C23Os degrade polycyclic aromatic hydrocarbons (PAHs) such as benzene, toluene, ethylbenzene, and xylenes, as well the degreaser and common groundwater contaminant trichloroethylene (TCE). Toluene has been identified among the organic compounds found in the humidity condensate samples from the US Space Shuttle Cabin [36]. The Shuttle did not use a water reclamation system, but the ISS does, as it reclaims water from urine and urine flush water, humidity condensate, personal hygiene water, and effluent from the crew health care systems. On a 1994 Shuttle mission, bags used to store urine samples for a life sciences experiment were giving off strong odors, and a post-flight assessment attributed this to the presence of volatile microbial metabolites which included 1,1,1-trichloroethane, and toluene [37]. In addition, trichloroethene (TCE) was found in one of two samples processed from the galley cold-water ports on Mir-21 [38] in the range of 1.8-2.3 ug/L – the EPA has set a maximum contaminant level (MCL) of 5µg/L (5 ppb) in drinking water for TCE (U.S. Environmental Protection Agency 1985). Therefore, in the event that trace amounts of PAHs are present in the PWS, *Burkholderia* species maybe using C23Os to catabolize the compounds into usable carbon sources in order to facilitate their survival in the low-nutrient environment of the PWS of the ISS.

Furthermore the observation that transposase-related genes are present at a mean copy number of 17.57 ± 5.7 per putative-plasmid, along with an integrase element with mean copy number of 2.47 ± 0.51, provides a possible beneficial mechanism when the population is under stress via promoting DNA rearrangement and duplications [36].

### *Burkholderia contaminans* plasmid encoded features

The coverage-based approach of plasmidSPAdes [40] identified putative plasmids in 2 of the *B. contaminans* isolates (s47 and s52). Both possess annotated T4SS and T2SS components as well as a soluble lytic murein transglycosylase harboring a putative invasion domain LysM at a mean copy number of 3 ± 2.83. The lytic transglycosylase, LtgG, has recently reported for its role to control cell morphology in *Burkholderia pseudomallei* and its virulence in the BALB-C mouse model [25]. Both also harbor a copy of the toxin component of the MazEF toxin-antitoxin system suggesting the ability for these ISS *B. contaminans* isolates to control their transition to the dormant persister state. This is because toxin-antitoxin systems are known to contribute to cell dormancy by halting cellular machinery through use of the toxin that is counteracted by the antitoxin once the stressor is removed [26]. Again, we see duplicated transposase-related genes with a mean copy number of 5 ± 5.7.

### Phenotypic assessment of ISS *Burkholderia* species

In order to test our hypothesis that these strains have the capacity to be virulent due to their plasmid gene content, we screened the ISS-derived *B. cepacia* and *B. contaminans* for the ability to invade and colonize macrophage. The *B. cepacia* isolates were found to invade macrophage by 6 hours, with some (s33) being more effective colonizers than others. They appear to multiply, then escape from the macrophage by 12 hours post-inoculation, possibly using the plasmid-encoded lysophospholipase mechanism to rupture the macrophage. Despite not contributing to a longer-term infection, the cytotoxic byproducts generated by lysophospholipase degradation of macrophage may play a physiological role in further stimulating the adhesion and differentiation of lymphoid cells macrophages and activation and recruitment of additional macrophage and T-lymphocytes, among other immune response mechanisms [24]. This is in contrast to the *B. cepacia* ATCC25416 terrestrial control strain, which has been noted for its ability to invade and carry out a long-term colonization of macrophage which can lend to the formation of long-term chronic infections [37]. We find that this strain displays the ability to remain intracellularized at 12-hours post-inoculation (Fig 5) yet remains able to lyse macrophage (Fig 4). Accordingly, the *B. cepacia* ATCC25416 reference strain contains additional virulence factors within its genome not found in the ISS isolates such as elements of the Type 6 secretion system (T6SS). *B. contaminans* isolates, on the other hand, seem to exhibit a decreased rate of cellular invasion and a more immediate cell lysis of macrophage. This follows suit with the hemolytic and antifungal capabilities they also display, which is not characteristic of the *B. cepacia* isolates. These ISS isolates appear slightly more virulent (as measured by LDH concentrations) than the incorporated *B. contaminans* terrestrial strain J2956, a pathogenic isolate from a sheep with mastitis. However, overall, the ISS *B. cepacia* and ISS *B. contaminans* isolates all appear to be less virulent to macrophage than the terrestrial reference *B. cepacia* strain.

## Methods

### DNA isolation

Bacterial isolates from the ISS PWD were obtained by filtering sample from both the ambient and hot water outlets of the potable water system through a microbial capture device (MCD) on the ISS by resident astronauts. The MCD filter was then placed on R2A agar and the plates were sent back to Earth for further isolation at JSC. Additionally, 1 liter of water was sent back in to Earth for further isolation at JSC on R2A agar. All isolates were cataloged and then stored at −80C in 15% glycerol until regrown on R2A agar slants and sent to JCVI. Upon arrival at JCVI all isolates were further plated on tryptic soy agar (TSA) and incubated overnight at 35°C, then inoculated into 4mls of tryptic soy broth (TSB). 2mls were stored with 15% glycerol as 1ml bacterial stocks and 2mls were centrifuged to form pellets with supernatant decanted then immediately frozen at −80C for future DNA extraction. DNA was extracted using a standard phenol/chloroform protocol. DNA was quantified using nanodrop and run on a gel to ensure high molecular weight samples were obtained.

### Genomic library preparation and sequencing

400ng of high molecular weight bacterial in 13ul of 1X TE (10mM Tris pH 8.0, 1mM EDTA) was processed using the NEBNext Ultra II FS DNA Library Prep Kit for Illumina (New England Biolabs, Ipswich, MA) protocol at half the standard reaction volumes. To the 13ul DNA, ul of NEBNext Ultra II FS Reaction Buffer and 1ul of NEBNext Ultra II FS Enzyme was added in a PCR tube and then incubated in a thermocycler at 37°C for 10 minutes at 37°C for fragmentation to target size, the enzyme is then denatured by incubating the reaction at 65°C for 30 minutes with lid set to 75°C. Next, adapter ligation is carried out on the 17.5 ul mixture of fragmented DNA by adding 15 ul of NEBNext Ultra II Ligation Master Mix, 0.5 ul of NEBNext Ligation Enhancer, 1.25 ul of NEBNext Adapter for Illumina for a total volume of 34.25 ul. The 34.25 ul adapter mixture is incubated at 20°C for 15 minutes a thermocycler with the heated lid off. This is followed by the addition of 1.5ul of USER Enzyme to the mixture, then incubation in the thermocycler at 37C for 15 minutes with heated lit set to 47C. In order to obtain 700-900 bp inserts, we used SPRIselect beads for a rightside clean-up of 0.25X, where the supernatant is retained for a left side clean up using a 0.25X bead clean up then eluted in 7.5 ul of 0.1X TE buffer. The fragmented, adapter ligated and size selected libraries were then amplified using 12.5 ul of NEBNext Ultra II Q5 Master Mix with 2.5 ul i7 index primer and 2.5 ul i5 Universal PCR primer for a total volume of 25 ul. The amplification was carried out at the following temperatures and times: initial denaturation at 98°C for 30 seconds, followed by 4 cycles of denaturation at 98°C for 10 seconds and annealing/extension at 65°C for 75 seconds, and a final extension at 65°C for 5 minutes. A total of 58 libraries were generated, quality controlled to find library size using the Agilent Bioanalyzer and high sensitivity DNA chip and double stranded DNA concentration was quantified using Qubit Fluorometric Quantitation. Libraries were normalized to achieve a total of 700 pM of pooled library in 200 ul with an average library size of 750 bps. An average of 7 million, 150 bp reads were obtained for the combined read 1 and read 2 of each library using Illumina’s NextSeq 500 High Output Kit.

### Genomic sequencing postprocessing

For the 24 libraries, demultiplexing was carried out allowing 1 bp mismatch and adapters were trimmed. bcl2fastq2 Conversion Software v2.17 was used to check for adapters, if detected, base calls matching the adapter and beyond the match are masked or removed in the resultant FASTQ file. Sequence quality was assessed using the program FastQC [38]. Genomic DNA library forward and reverse reads were quality trimmed using trimmomatic v0.39 [39] with a sliding window of 5:20 and a minimum length of 100 base pairs. Libraries were de novo assembled using SPAdes v3.12.0 [14]. Assembly summaries are presented in S2 Table. Average nucleotide identity was calculated with fastANI v1.2 [15]. An estimated maximum-likelihood based with GToTree v1.4.4 [16] based on amino acid sequences of 203 single-copy genes specific to Betaproteobacteria using FastTree2 v2.1.10 [40]. Whole-genome assembly single-nucleotide variant (SNV) trees were generated with Parsnp v1.2 [17].

All pangenomic analyses were performed within anvi’o v5.5 [19], which uses MCL clustering [18]. Default settings were used other than setting the ‘--mcl-inflation’ parameter to 7.

### Functional enrichment/depletion analysis

This was performed within anvi’o v5.5 (citation) utilizing the ‘anvi-get-enriched-functions-per-pan-group’ program with default settings. The resulting table was filtered to include only those with a Benjamini-Hochgerg corrected p-values of <= 0.1.

### Macrophage infection

The infection assay aimed for a multiplicity of infection (MOI) of 5 ISS *B. cepacia* and *B. contaminans* cells for every one macrophage cell. Biological replicates of the experiment were conducted on three separate days to ensure a robust reproducibility in results. At time zero CFU/mL count was conducted for inoculum of each ISS isolate and was averaged for each biological replicate experiment of the separate days (S1 Fig). These counts showed that our experimental MOI was in the range of MOI 5-11 with an average of MOI of 7 over the three replicated experiments. The bacteria were allowed to infect the macrophage cells for two hours before being washed away and replaced with fresh media containing 15ug/mL ceftazidime and 1mg/ml amikacin to ensure clearance of extra cellular bacteria. At six, eight, twelve and 24 hours post-infection, the ISS treated sample supernatant and control supernatant from uninfected cells is collected and macrophage cells were removed by centrifugation at 300rpm for 2mins. The supernatant was then assessed for the presence of lactase dehydrogenase (LDH) using a Cytotoxicity Detection Kit (LDH) (Roche, Germany). The cell-free supernatant is incubated with the reaction mixture from the kit. LDH activity is determined using an enzymatic test. In the first step NAD+ is reduced to NADH/H+ by the LDH-catalyzed conversion oflactate to pyruvate. In the second step the catalyst (diaphorase) transfers H/H+ from NADH/H+ to the tetrazolium salt INT which is reduced to formazan. The enzyme reaction was stopped by the addition of 50 l/ well 1N HCl (final concentration: 0.2 N HCl) after 10 minutes of incubation. The formazan dye absorbance was measured using the 490nm wavelength. We present the LDH data for all ISS strains and *B. contaminans* J2956 and *B. cepacia* ATCC25416 reference strains (S2 Fig, all LDH assay panels). Additionally, the macrophages were washed after removal of the supernatants at six, eight, twelve- and 24-hours post-infection to remove any dead extracellular bacterial cells, then lysed in order enumerate the intracellular bacterial cells by plating and colony counting.

### Aspergillus fumigatus screen

JCVI −80°C freezer stocks were streaked onto TSA and incubated 48 hours at 37°C, single colonies were then inoculated into 3 mLs TSB and incubated overnight at 37°C with shaking at 220rpm. Aspergillus fumigatus AF293 strain was maintained as glycerol stocks and grown at 37°C. The fungus was incubated on Potato Dextrose Agar (PDA; Acumedia Manufactures Inc. Lansing, MI) at 37°C under the dark for 4 days. The fungal spores were collected with loops and suspended with 0.01% tween 80 sterilized water. The spore suspension was counted by hemocytometer and was inoculated to 10^6 spores per plate to PDA and TSA media, concurrently 5 μl of a *Burkholderia* isolate was spotted in a quadrant of each of the agar types. The samples were incubated at 37°C and the zones of inhibition were recorded after 48hours.

### Hemolysis assay

JCVI −80°C freezer stocks were streaked onto TSA and incubated 48 hours at 37°C, single colonies were then inoculated into 3 mLs TSB and incubated overnight at 37°C with shaking at 220 rpm. 5 μl of each *Burkholderia* isolate culture was spotted onto a petri plate (100mm × 15mm) containing TSA with 5% sheep’s blood and incubated at 37°C for 48 hours.

### MIC determination

1xMICs of all antibiotics were determined using the National Committee for Clinical Laboratory Standards (NCCLS). In the MIC assay, 2ul of an antibiotic dilution series (beginning at 64ug/mL) was used to treat 100ul of OD_600_=0.001 culture (diluted from cells in exponential growth phase), then incubated without shaking for 18 hours before MIC determination. The MIC value is the lowest concentration at which bacterial growth is fully arrested.

### Biofilm assay

JCVI −80°C freezer stocks were streaked onto TSA and incubated 48 hours at 35°C, single colonies were then inoculated into 3 mLs TSB and incubated overnight at 3°C with shaking at 220 rpm. Cultures were diluted to an OD600 of 1, then diluted 1:100, each culture was then plated into one column of three 96 well polystyrene plates at 150 μl per well. The cultures were covered with adhesive film, then incubated at 35°C for 24, 48, and 72 hours. At each timepoint an entire 96 well plate having 11 cultures and control culture were stain using crystal violet. This process includes removing the cultured media from each well by inverting the plate and giving it 3-5 abrupt shakes into a waste container, then rinsing the wells lightly with water. This is followed by the addition of 150 μl of crystal violet into each well, the crystal violet is incubated in the well to adhere to any remaining biofilm for exactly 15 minutes, then the plate is carefully inverted and given 3-5 abrupt shakes into a crystal violet waste container. The plate is carefully rinsed with water and then laid upside down to dry. Once each of the 3 time points have been processed, the crystal violet is resolubilized for each timepoint plate using 100% EtOH by pipette mixing, then transferred to a new plate and covered to avoid evaporation. The resolubilized crystal violet strain is then placed in a plate reader to obtain an absorbance at 590 nm.

## Data availability

The genomes reported in this paper can be found under bioproject: PRJNA493516.

## Acknowledgments

AOR was funded by NASA Space Biology under grant 80NSSC17K0035.

MDL was funded by NASA Space Biology under grant NNH16ZTT001N-MOBE.

CLD was funded by the NASA Astrobiology Institute Alternative Earth’s.

## Author Contributions

Experiments were conducted by AOR. Bioinformatic analyses were conducted by AOR, MDL, and CLD. The manuscript was written by AOR and ML and CLD. Experiments were devised in collaborative effort by AOR, WCN, and CLD.

## Supplemental Figures

**Supplemental Figure1:**
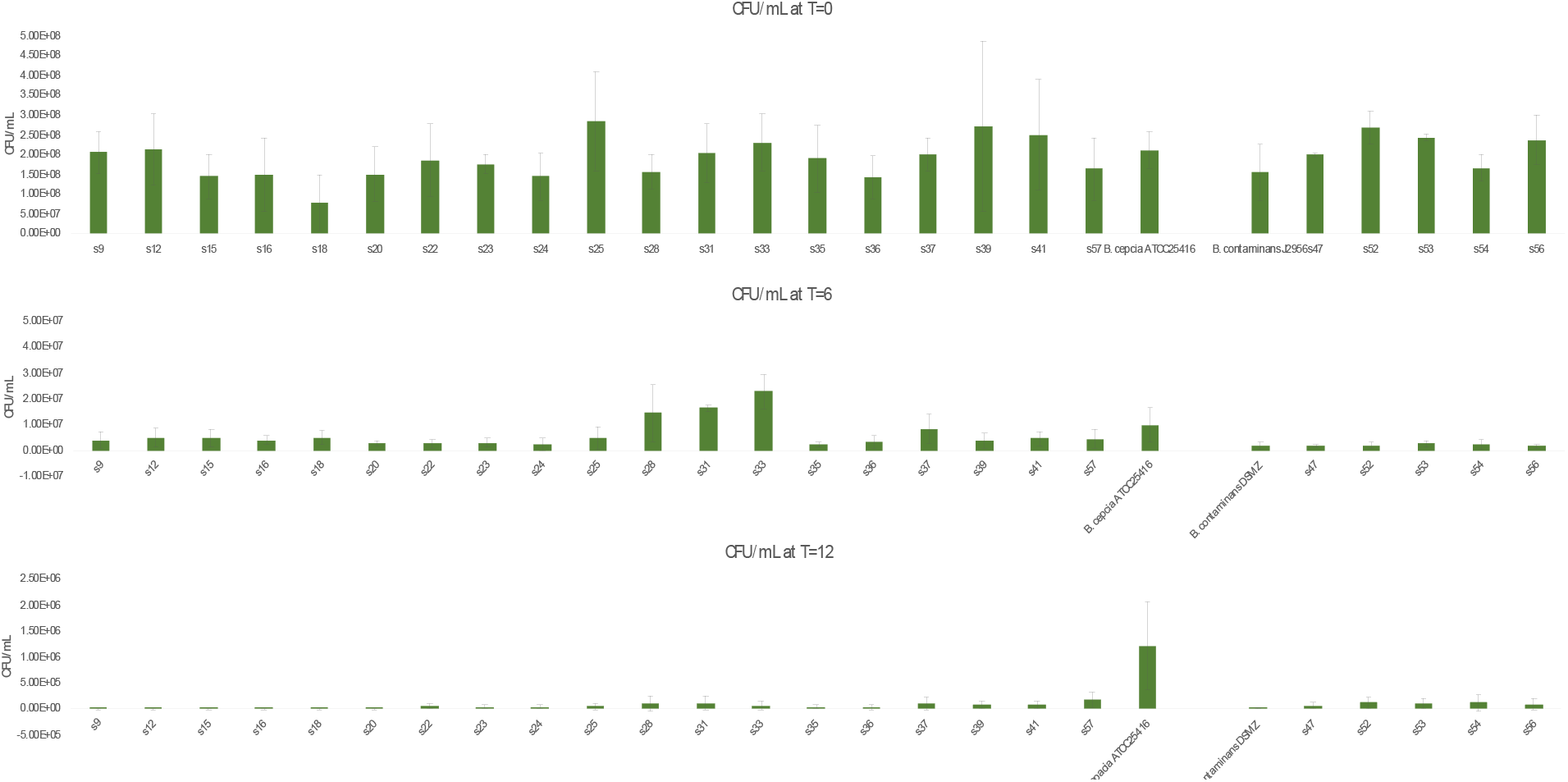
Macrophage experiment CFU/mL time course data.

**S2 Fig.**
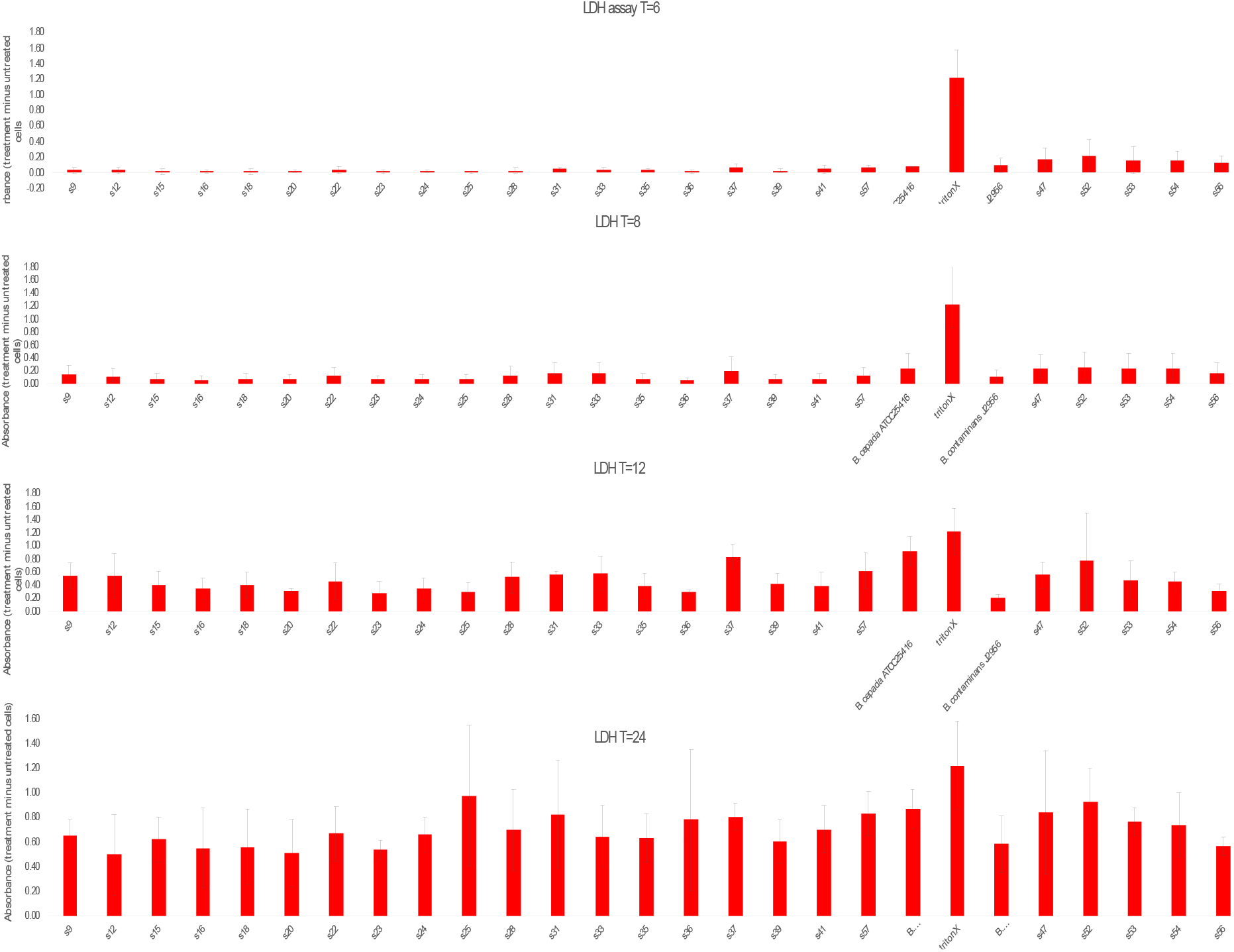
Macrophage experiment LDH time course data.

## Supplemental Tables

**S1 Table: Dates of isolation and sample number to NASA identifier.**

**S2 Table: Spades Assembly data**

**S3 Table: B. contaminans functional enrichments**

**S4 Table: B. cepacia functional enrichments**

**S5 Table: ISS isolate only pangenome with annotations and sequences**

**S6 Table: ISS isolate only plasmid pangenome with annotations and sequences**

